# Sleeve gastrectomy promotes sustained weight loss while increasing heat production in middle aged, obese female mice

**DOI:** 10.1101/2020.10.04.325787

**Authors:** Ana BF Emiliano, Ying He, Sei Higuchi, Rabih Nemr, Natalie Lopatinsky, Gary J. Schwartz

## Abstract

**Background:** Sleeve gastrectomy (SG) is currently the most frequently performed bariatric surgery in the United States. The majority of patients undergoing SG are middle aged women. Most preclinical models of bariatric surgery, however, utilize juvenile male mice. A long-term characterization of the response of mature wild type, obese female mice to SG has not been performed. Thus, we set out to characterize the response of middle aged obese female mice to SG.

**Methods:** Ten-month old C57bl/6J obese female mice were randomized to undergo SG, sham surgery without caloric restriction (SH) or sham surgery with caloric restriction to match body weight to the SG group (SWM). Body weight, body composition and glucose tolerance were matched at baseline. Mice were followed for 60 days following their respective surgeries.

**Results:** The SG group had a more pronounced percent weight loss than the SH and SWM control groups (p<0.05), while consuming more calories than the SWM group (p<0.05). The SG group had a significant improvement in glucose tolerance compared to the SH control group (p<0.05). Plasma leptin was significantly decreased in the SG and SWM group, compared to the SH group (p<0.01). Unexpectedly, FGF-21 was increased in the SH group compared to the SG and SWM groups (p<0.01), while there was no difference in plasma insulin among the three groups. Heat production was increased in the SG group compared to SWM and SH groups (p<0.001). SG also had a significantly increased mRNA expression of Uncoupling Protein 1 (UCP-1), Adiponectin and Peroxisome Proliferator-Activated Receptor Gamma Coactivator 1-alpha (PGC1-alpha) in brown adipose tissue (BAT), compared to SWM and SH groups. Both SG and SWM groups had increased fecal lipid excretion (p<0.05), compared to the SH group.

**Conclusions:** SG in obese, middle aged female mice leads to sustained weight loss and blood glucose improvement. It appears that increased metabolism in BAT may be linked to these effects.

## Introduction

Obesity and type 2 diabetes are highly prevalent disorders, affecting millions of Americans^1^. Due to the limited efficacy of medical therapies to decrease mortality associated with these conditions, there has been an increased demand for bariatric surgery^2^. To date, bariatric surgeries including Sleeve Gastrectomy (SG) and Roux-En-Y Gastric Bypass (RYGB) have higher efficacy in promoting durable weight loss and lowering blood glucose in patients with obesity and type 2 diabetes than medical therapies alone^3,4,5^. Currently, SG is the most frequently performed bariatric surgery in the US, at a rate of 60% of all bariatric procedures^6^. Bariatric surgery studies indicate that most patients seeking and undergoing bariatric surgery are middle-aged women^7,8^. In the quest to understand the mechanistic underpinnings of the sustained weight loss and glycemic improvement secondary to bariatric surgery, rodent models have become invaluable tools^9,10,11^. However, one of the weaknesses of these same models is the almost exclusive utilization of juvenile male mice, with a few exceptions that, in spite of using females, still fail to use more mature mice in their studies^12,13^. Although information obtained from these preclinical rodent models can be extremely useful, they are also limited when these models do not reflect the human patient population that the study is targeting, whether adolescents, young adults, middle aged or older adults.

C57bl6/J wild type mice is a common strain used in obesity and hyperglycemia studies, since these mice gain weight when fed a high fat diet (diet-induced obese or DIO model)^14^. Most transgenic and knock-in mouse models are bred in a C57bl6/J background, as well. However, obese female C57bl6/J are not commercially available. This limits attempts to compare studies using C57bl6/J females to studies performed using male DIO mice, if the obese female mice are from a different strain. Another important issue is the age of the mice, since it is not clear if younger mice have a better response to bariatric surgery than older, more mature mice. The average age of the typical bariatric surgery patient is between 45-50 years old, which is definitely not the age range of mice in most rodent bariatric studies. Moreover, the chronicity of exposure to high fat diet may also be significant for the response to bariatric surgery. It is possible that mice that are fed a high fat diet for 8 weeks cannot be compared to mice fed a high fat diet for 16 weeks or longer. Intuitively, one would tend to think that a shorter exposure to high fat diet would lead to faster weight loss and more significant improvement in blood glucose, than a longer and more chronic exposure. It is important to note that human patients with obesity are chronically exposed to the conditions that lead to obesity. Most individuals with obesity have been obese for life.

The present study was designed to tackle the issues discussed above. We made our own C57bl/6J female DIO, starting the mice on a 60% high fat diet at 6 weeks of age. When they reached middle age, at approximately 10 months old, these mice were randomized into a SG group, sham surgery with ad libitum diet (SH) or sham surgery weight-matched (SWM) to the SG group, through caloric restriction. We found that these older, mature obese mice responded well to SG, displaying durable weight loss and glycemic improvement. Although by the end of the study they had a similar body weight as the SWM group, their caloric intake was higher. Our findings point to increased energy expenditure through heat production as a possible mechanism underlying their significant and durable weight loss.

## Methods

### Animals

All experiments were performed in accordance to the Columbia University Institutional Animal Care and Use Committee. Female C57bl/6J were started on a high fat diet at 6 weeks of age (Research Diets, catalogue number D12492, New Jersey, NJ). At 10 months of age, mice were randomized to SG, SH and SWM to the SG group (n=13 for SG; n=6 SH, n=6 SWM). Mice were single-housed. Mice were sacked after 60 days post-operatively, after a 5 hour fast, under isoflurane anesthesia, when intracardiac blood collection and tissue harvest were performed. Body weight and food intake were measured daily for the first week post-operatively and then weekly. Body composition with EchoMRI was done at baseline, at one month and two months post-operatively. Glucose tolerance tests were done at baseline, and then at one, four, six, eight and ten weeks post-operatively. Feces were collected for fecal lipid content two months post-operatively. Plasma and tissue samples for mRNA analysis were also obtained 60 days post-operatively. Mice spent one week in metabolic cages between 7-8 weeks post-operatively. Baseline measurements are defined as having being taken 2 weeks prior to surgeries.

### Sleeve gastrectomy technique

All surgical procedures were sterile. Mice received meloxicam, saline and enrofloxacin subcutaneously at the time of the surgery. A midline laparotomy was performed and the stomach isolated from surrounding connective tissue. Surgeries were performed under a dissecting microscope (Leica M-125). For SG, the gastric arteries were ligated with 8-0 suture (Vicryl Violet J401G, BV130-5 Taper, Ethicon). After ligation, an incision was made at the bottom of the fundus and the stomach contents emptied. The stomach was irrigated with warm saline. A 6-0 suture (Vicryl J212H, Ethicon) was used for continuous suturing from about 2 millimeters below the esophagogastric junction, to the point where the pancreas is attached to the stomach. Tissue below that line was dissected with microscissors. The edges of the stomach were sutured together with 7-0 suture (Vicryl J488G, P-1 cutting, Ethicon). Gastric leaks were assessed. Sham surgeries were as above, except that after stomach isolation, the stomach was put back into the abdominal cavity and the abdominal wall closed in layers with 5-0 suture (Vicryl J493G, Ethicon). Mice were kept on a liquid diet with Ensure High Protein (Abbott Laboratories) and high fat diet for 4 days prior to the surgeries. The day before the surgery, high fat diet was removed and the mice were kept on Ensure until the 7^th^ post-operative day. High fat diet was reintroduced on the 5^th^ post-operative day and continued until the end of the study. SG and Sham groups were kept on ad libitum high fat diet. The sham weight-matched group was kept on 2-2.5 grams of high fat diet daily, provided at different times of the day to prevent entrainment. The SG surgery described here is a modification of an SG surgical technique previously developed by another group^15^.

### Glucose Tolerance Test

Oral glucose tolerance test (OGTT) was performed by oral gavage using a plastic needle (1FTP-20-38, Instech Laboratories, Inc.), with 2 grams of dextrose per kg of body weight. Intraperitoneal glucose tolerance test was performed by injecting the mice intraperitoneally with 2 grams of dextrose per kg of body weight.

### Energy expenditure

Mice were in Comprehensive Lab Animal Monitoring System (CLAMS, Columbus Instruments, Columbus, OH) for one week, between 7-8 weeks post-operatively.

### Body Composition

Body composition was measured at baseline, at one and two months post-operatively using Echo-MRI^™^-100H (EchoMRI LLC, Houston, TX).

### Fecal Lipid Content Assay

Feces were collected from individual mice. Feces were dried overnight on a six-well plate, at 42°C. For each reaction, 100 mg of feces (per mouse) was placed in 1 mL of NaCl and then homogenized with beads for five minutes. The solution was transferred to a 15 mL tube, to which a mix of chloroform and methanol at 2:1 was added (Chloroform, Fisher, C298-4; Methanol, Fisher A456-212). The mixture was vortexed vigorously and then centrifuged at 2000g for 10 minutes. The chloroform phase was collected and transferred to a 20 mL vial and evaporated under N2. After that, 1 mL of deionized water was added. The next day, samples were assayed in duplicates, with a Free Fatty Acid (FFA) kit (Wako 995-34693; 997-34893; Fujifilm USA). Samples were diluted at 1:20 and incubated at 37°C for five minutes and read with a microplate reader.

### RNA extraction and analysis

RNA extraction was performed using NucleoSpin RNA Set for Nucleozol (Macherey-Nagel, catalogue #740406.50). cDNA was obtained using the High Capacity cDNA Reverse Transcription Kit (Applied BioSystems, ThermoFisher Scientific, catalogue #4368813). For RT-PCR, we used ready-made probes purchased from Integrated DNA Technologies (IDT, USA). The enzyme used was Taqman Fast Advanced Master Mix (ThermoFisher Scientific, catalogue #4444557). Samples were run in triplicates and the experiment replicated three times. The instrument was QuantiStudio^™^ 5 Real Time PCR system (ThermoFisher Scientific). We used the QuantiStudio^™^ 5 software to perform the analysis.

### Plasma Assays

Blood was collected from animals fasted for 5 hours, via intracardiac puncture. Blood was collected in tubes containing EDTA, aprotinin and Diprotin A, a DPPIV inhibitor (EDTA, ThermoFisher Scientific, catalogue #15575020; Aprotinin, Millipore Sigma, catalogue #A1153; Diprotin A, Enzo Scientific, catalogue #ALX-260-036-M005). Blood was spun for 15 min at 2000 g, in a refrigerated benchtop microcentrifuge and subsequently stored in a minus 80 freezer. For FGF-21, Leptin and Insulin plasma measurement, we used a U-PLEX MSD assay (MesoScaleDiscovery, catalogue #K152ACL-1).

### Estrus cycle

We performed vaginal smears from all the mice in the study, to assess whether they were still having estrus cycles, considering their age. We collected 30 microliters of saline that was gently introduced in the vaginal canal and then proceed with microscopic analysis using under bright field illumination (Nikon microscope, Eclipse series).

### Statistical Analysis

We used GraphPad Prism 8.0 version software for all statistical analysis. One Way ANOVA was used for comparison of three groups. Two-Way ANOVA with repeated measures was used for comparisons of three groups with multiple measures. XY analysis was used for generating AUC graphs used for glucose tolerance and energy expenditure analysis, which were subsequently analyzed with ANOVA. Power was set at 80%, with a p value of <0.05.

## Results

### Body Weight

Mice at baseline had body weight, fat mass and lean mass that were not statistically different, with mice having an average body weight of approximately 40 grams (figure 1, top panel). From our experience, female C57bl/6J do not gain as much weight as male C57bl/6J mice, in spite of having been maintained on a high fat diet for more than 30 weeks. One month after the surgery, both SG and SWM had statistically lower body weight and fat mass, compared to the sham group (p<0.05) (figure 1, middle panel). At two months, SG and SWM had even lower body weight and fat mass (P<0.01) and now they also had lower lean mass than the sham group (p<0.05) (figure 1, bottom panel). Percent weigh loss was not significant in the first month. However, it was significant in the second month post-operatively (p<0.05) (figure 3, month 1 and month 2). Fecal lipid content was higher in SG and SWM, than in the sham group, possibly indicating malabsorption (p<0.05) (figure 3, fecal lipid content).

**Figure 1.**
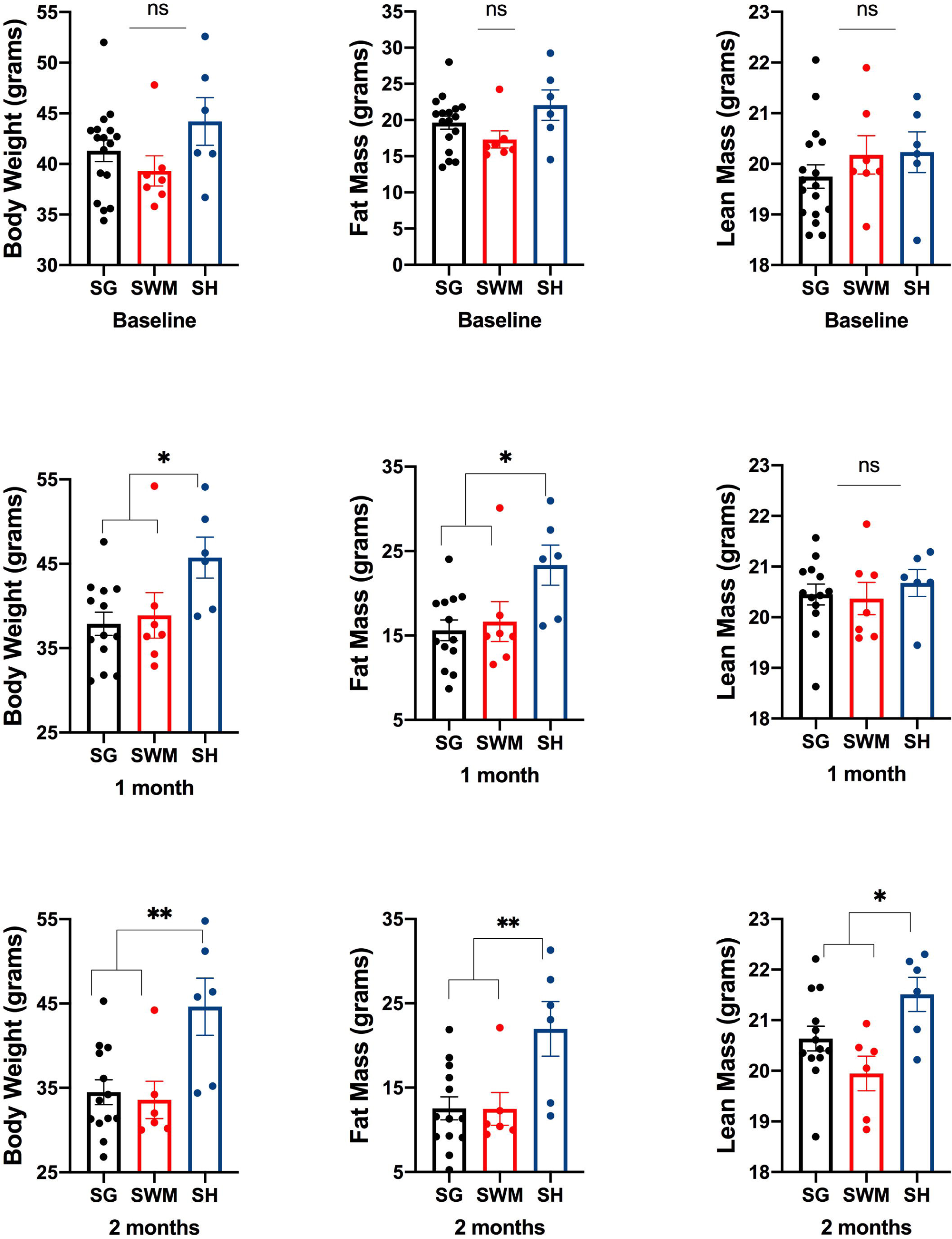
Top panel depicts body weight and body composition at baseline. Middle panel depicts body weight and body composition in the first month after surgeries. Bottom panel shows body weight and body composition in the second month after surgeries. SG – sleeve gastrectomy group; SWM – sham weight matched; SH – sham. NS – nonsignificant; * p<0.05; ** p<0.01. For n and statistical analysis, please see text.

**Figure 2.**
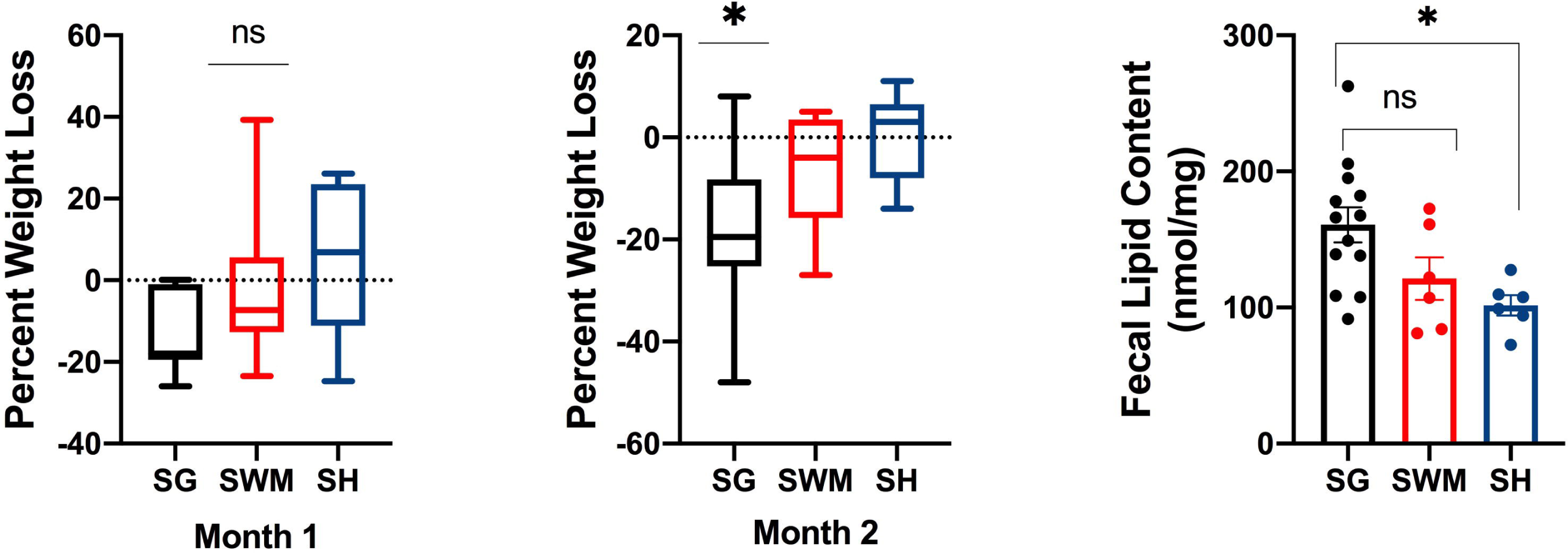
First two panels show percent weight loss at 1 and 2 months after surgeries. Last panel shows fecal lipid content. SG – sleeve gastrectomy group; SWM – sham weight matched; SH – sham. NS – nonsignificant; * p<0.05; ** p<0.01. For n and statistical analysis, please see text.

**Figure 3.**
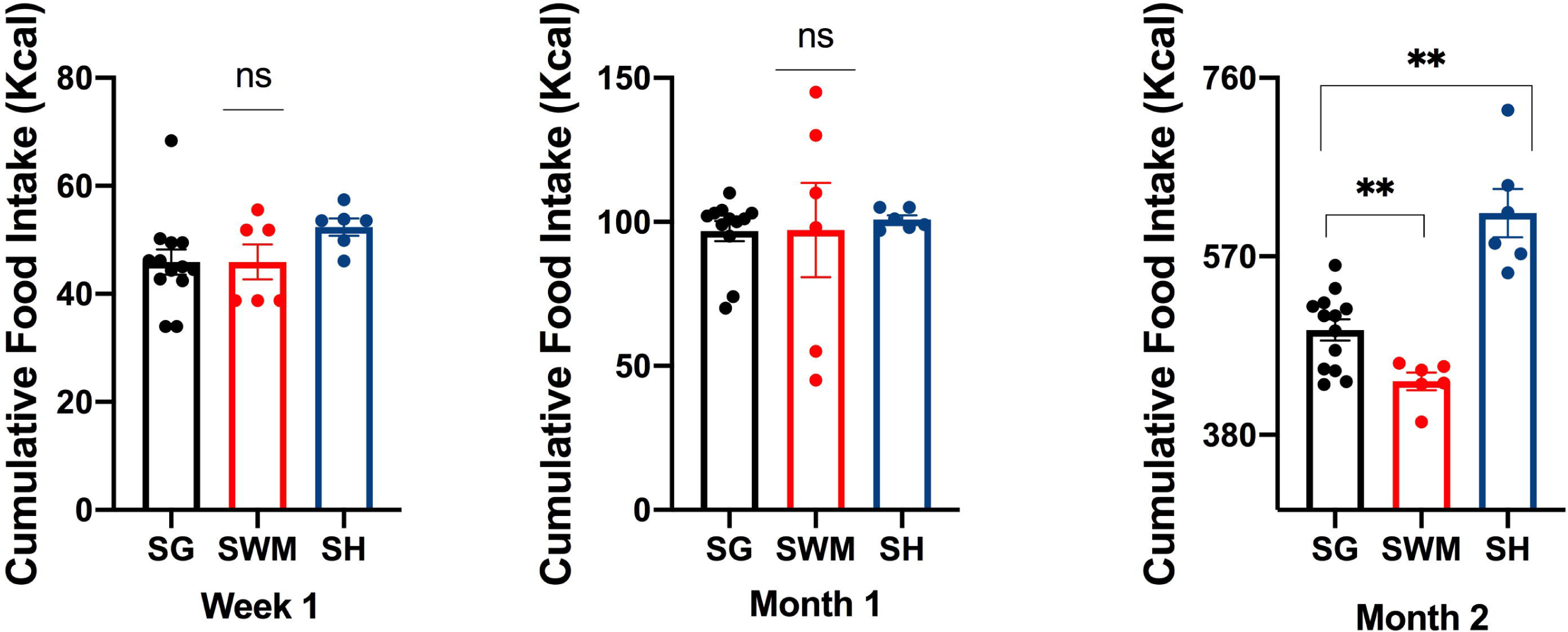
Cumulative food intake at the first post-operative week and then at 1 and 2 months after surgeries. SG – sleeve gastrectomy group; SWM – sham weight matched; SH – sham. NS – nonsignificant; * p<0.05; ** p<0.01. For n and statistical analysis, please see text.

### Food Intake

In spite of reports of lower food intake in the first two weeks post-operatively, based on male mouse studies, in the first week and first month, SG had the same food intake as the two other control groups (p>0.05) (figure 3, week 1 and Month 1). By the second month post-operatively, SG consumed significantly fewer calories than the sham group but consumed significantly more calories than the SWM group (p<0.01) (figure 3, Month 2).

### Glucose Tolerance

IPGTT at baseline did not reveal no difference in AUC among the three groups, as all the mice were severely hyperglycemic (p>0.05) (figure 4, top panel). OGTT done one week post-operatively, showed a significantly lower AUC for SG compared to both SWM and SH (p<0.001) (figure 4, middle panel). At one month, there was no longer any difference in AUC among the groups (p>0.05) (figure 4, bottom panel). At 6 weeks post-operatively, the IPGTT AUC was significantly lower in the SG group compared to the SWM and SH groups (p<0.05) (figure 5, top panel). At 8 weeks post-operatively, the OGTT AUC was significantly lower in the SG and SWM groups, compared to SH control (p<0.05) (figure 5, middle panel). At 10 weeks post-operatively, the IPGTT AUC was significantly lower in the SG and SWM groups, compared to SH control (p<0.05) (figure 5, bottom panel).

**Figure 4.**
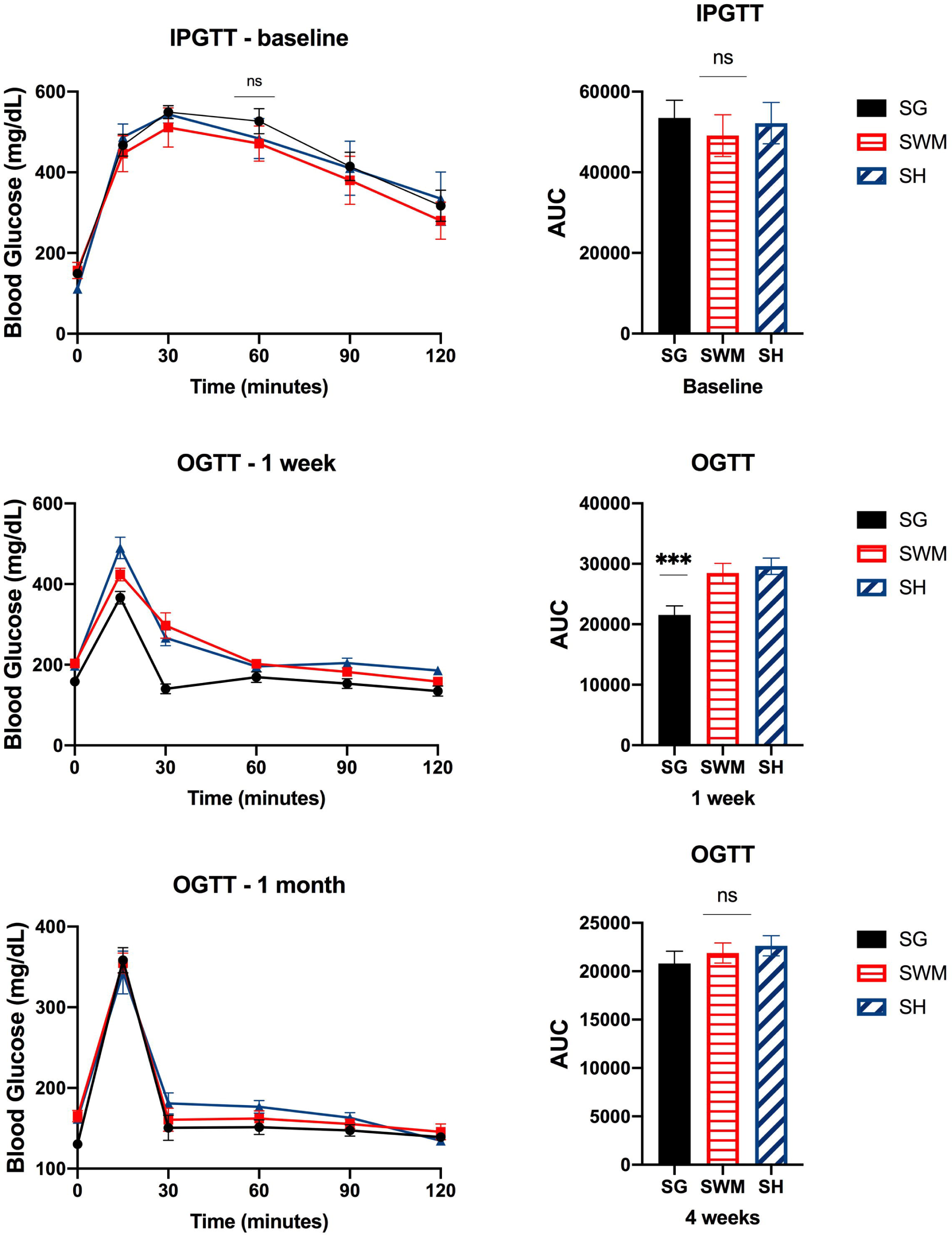
Top panel shows baseline IPGTT and IPGTT AUC. Middle panel shows OGTT and OGTT AUC at 1 week post-operatively. Bottom panel shows OGTT and OGTT AUC 1 month post-operatively. SG – sleeve gastrectomy group; SWM – sham weight matched; SH – sham. NS – nonsignificant; * p<0.05; ** p<0.01. For n and statistical analysis, please see text. IPGTT-intraperitoneal glucose tolerance test; OGTT – oral glucose tolerance test.

**Figure 5.**
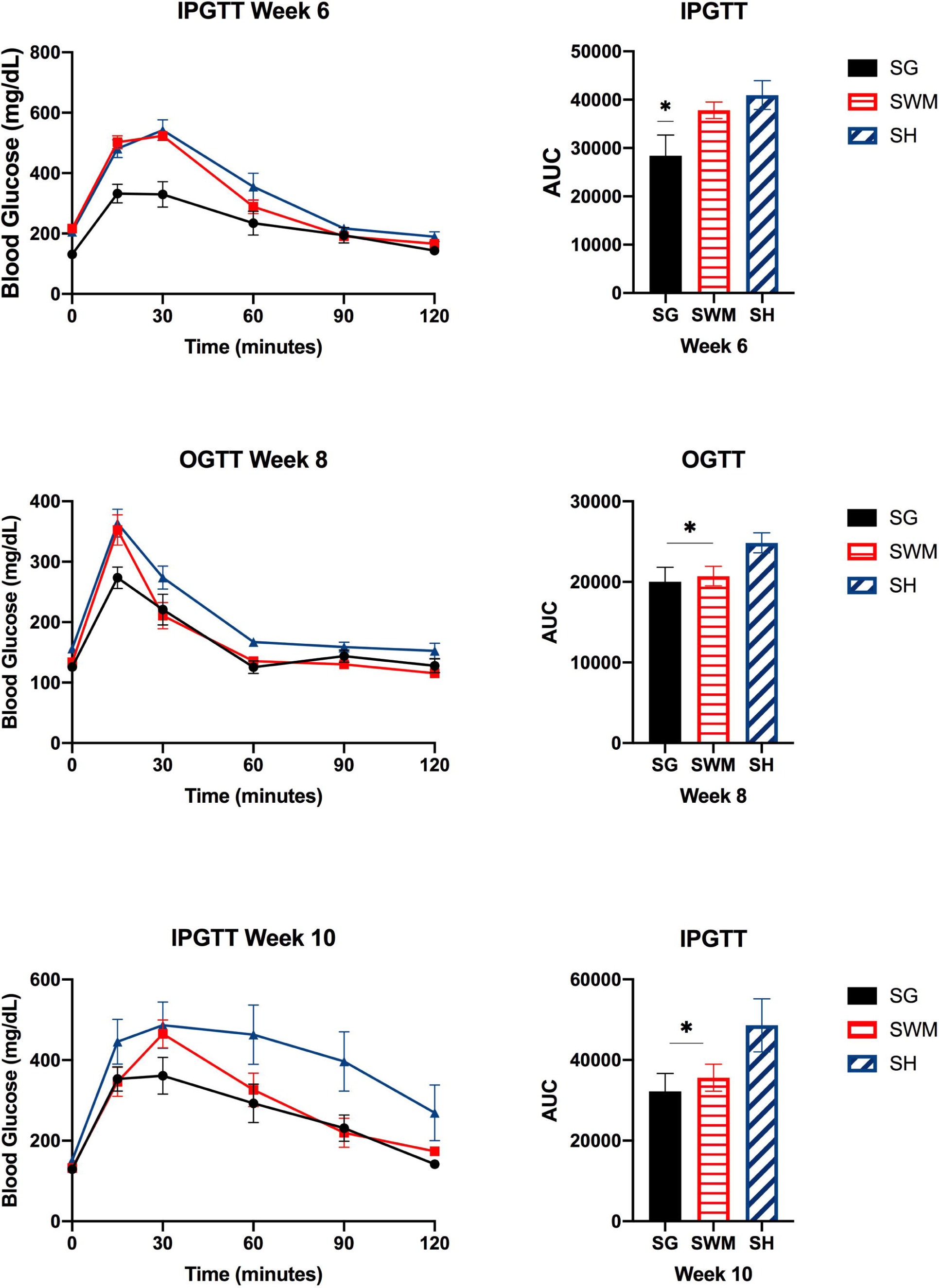
Top panel shows IPGTT and IPGTT AUC at 6 weeks post-operatively. Middle panel shows OGTT 8 weeks post-operatively. Bottom panel shows IPGTT 10 weeks post-operatively. SG – sleeve gastrectomy group; SWM – sham weight matched; SH – sham. NS – nonsignificant; * p<0.05; ** p<0.01. For n and statistical analysis, please see text. IPGTT-intraperitoneal glucose tolerance test; OGTT – oral glucose tolerance test.

### Plasma hormones

At the end of the second month post-operatively, plasma FGF-21 was significantly higher in the sham control group, compared to the SG and SWM groups (p<0.01) (figure 6). Plasma insulin was not different among the groups, in spite of a trend towards being lower in the SG and SWM groups (p=0.09). Plasma leptin was significantly lower in the SG and SWM groups, compared to the sham group (p<0.01).

**Figure 6.**
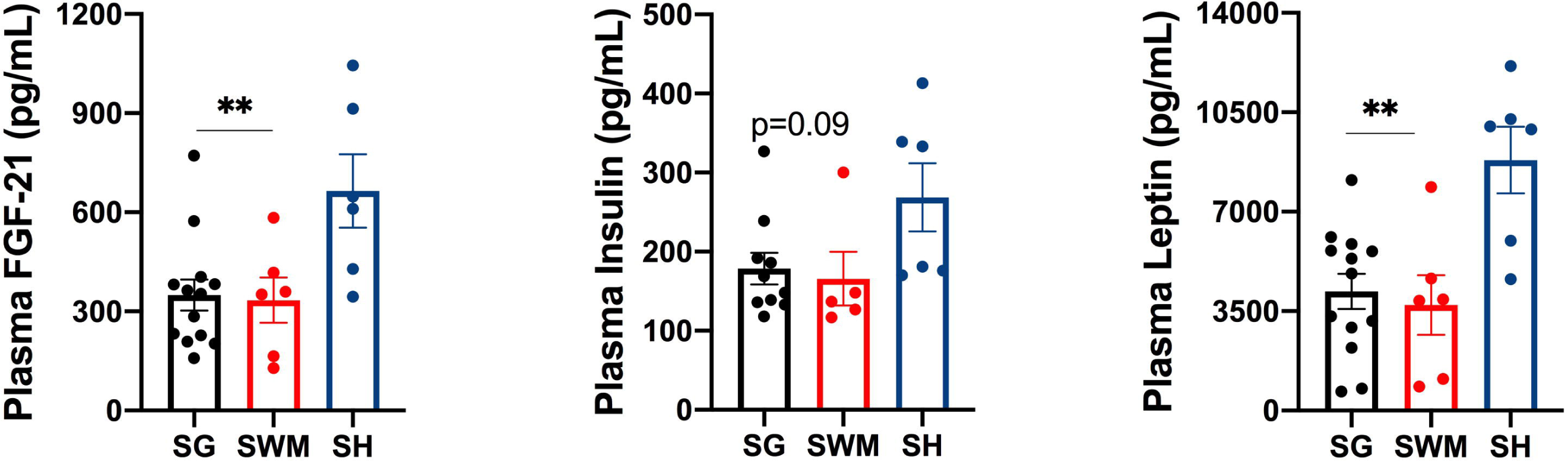
Plasma levels of FGF-21, insulin and leptin. SG – sleeve gastrectomy group; SWM – sham weight matched; SH – sham. NS – nonsignificant; * p<0.05; ** p<0.01. For n and statistical analysis, please see text.

### Estrus cycle

We collected evidence that the mice were in various phases of the estrus cycle, including metestrus, diestrus, proestrus, and estrus (figure 7).

**Figure 7.**
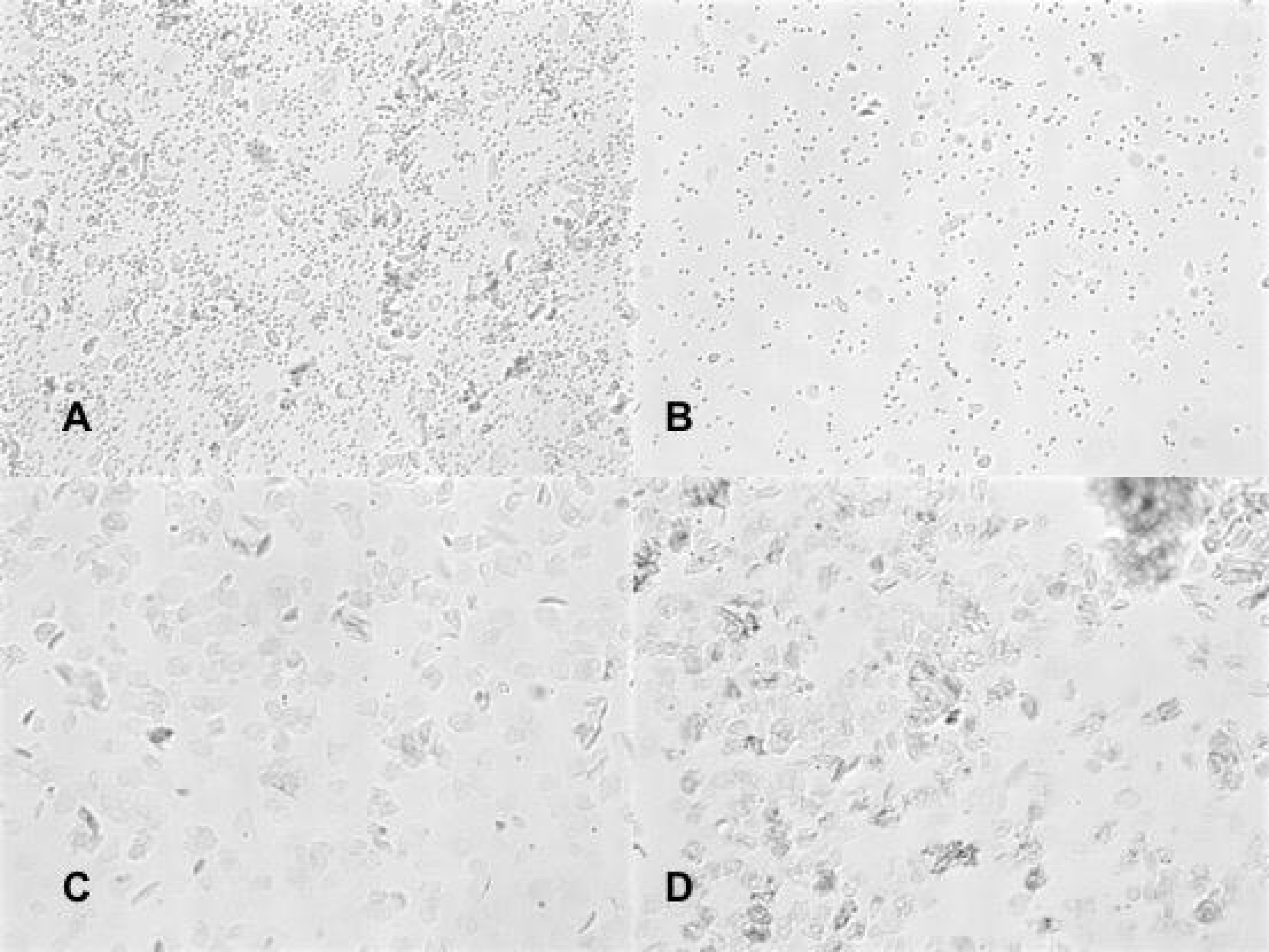
Estrus cycle phases as observed in the mice cohort used in all experiments. A – Metestrus; B-Diestrus; C-Proestrus; D – Estrus.

### Energy expenditure and BAT mRNA expression

The SG group had an increased mRNA expression of genes involved in metabolic activity in the BAT, namely Uncoupling Protein 1(UCP-1), Peroxisome proliferator-activated receptor gamma coactivator 1-alpha (PGC1 alpha) and Adiponectin (figure 8, top panel). The SG mice also displayed greater heat production than the other two control groups (p<0.001) (figure 8, middle panel). The Respirator Exchange Ratio (RER) was not different between SG and the sham group (p<0.05), but it was significantly lower in the SWM group compared to the other two groups (p<0.0001) (figure 8, bottom panel).

**Figure 8.**
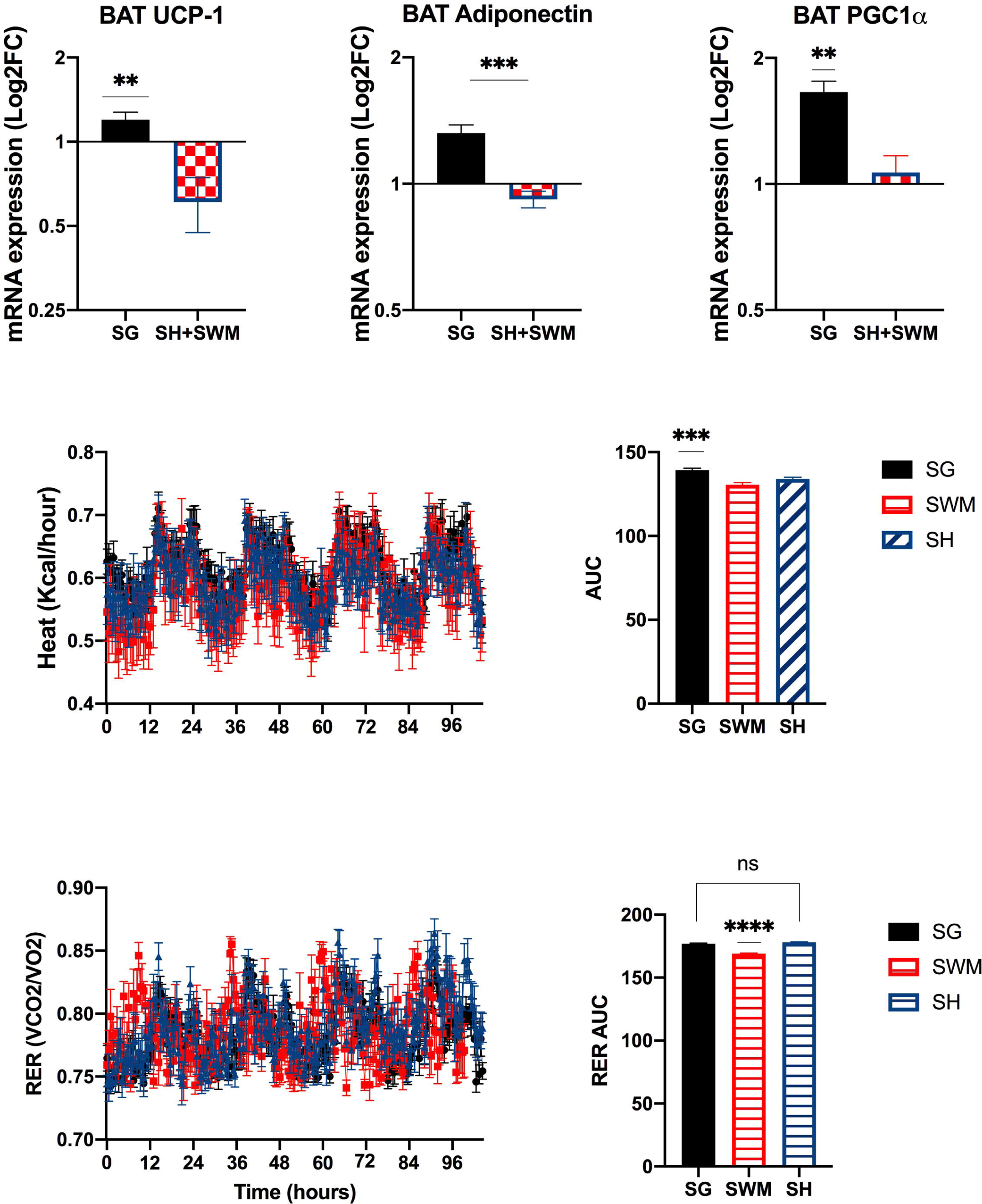
Top panel shows mRNA expression in BAT mice, including UCP-1, Adiponectin and PGC1-alpha. Middle panel shows heat production over time. Bottom panel shows RER over time. SG – sleeve gastrectomy group; SWM – sham weight matched; SH – sham. NS – nonsignificant; * p<0.05; ** p<0.01; ***p<0.001; ****p<0.0001. For n and statistical analysis, please see text.

## Discussion

Using mouse models that have similar phenotypic characteristics to the human patient population whose disease is being studied is fundamental to generate meaningful translational research. Obesity in adolescence and young adulthood has different characteristics than obesity in older age groups. Lifelong obesity has more sequelae because of the accrual of longstanding metabolic impairments started at a younger age^16^. In the present study, we used a mouse model of obese female mice corresponding in age to middle-aged obese women. We found that mature, obese female C57bl6/J mice are severely hyperglycemic at baseline but respond to SG with significant and sustained weight loss and blood glucose improvement compared to the SH control group. It was notable that the SG group achieved a similar weight compared to the SWM group, in spite of a higher caloric intake. One possibility would be malabsorption induced by SG, as suggested by the increased fecal lipid content found in the SG group. However, an increased fecal lipid content was also found in the SWM group. After sham surgery, mice have an intact stomach, as confirmed by our own necropsy studies. Therefore, we cannot ascribe malabsorption as the cause of increased fecal lipid content, either in the SG or SWM group. SG is not typically considered a malabsorptive bariatric procedure^17^. Gut microbiome changes secondary to weight loss have been associated with a decrease in intestinal fat absorption^18^. It is unclear whether this could be the reason for the increased fecal lipid content found in the SG and SWM groups.

Other factors that could account for the more pronounced weight loss in the SG are, the finding of a significantly increased heat production in the SG group, as well as increased BAT expression of UCP-1, adiponectin and PPARgc1-alpha mRNA in the SG group, compared to SWM and SH groups. BAT is the effector organ for non-shivering thermogenesis in rodents, and UCP-1 is key for that function. When activated by cold or dietary intake, UCP-1 uncouples mitochondrial fatty acid oxidation from the production of ATP, and the energy is dissipated as heat^19^. UCP-1 ablation induces obesity in mice, as long as their environment is thermoneutral^20^. However, increased UCP-1 expression per se in rodent models has not been shown to increase basal metabolism^21^. A stimulus leading to UCP-1 activation is also necessary for the UCP-1 increased expression to translate into increased basal metabolism. Accordingly, we also found a significantly increased expression of PGC1-alpha mRNA in BAT of SG mice. PGC1alpha transcription is increased by sympathetic activation and by exposure to cold temperatures, inducing nonshivering thermogenesis. PGC1alpha coactivates nuclear receptors that bind to the UCP-1 enhancer^22^. Furthermore, we also found that adiponectin mRNA expression is increased in the BAT of SG mice, compared to the control groups. Adiponectin knock-out mice fail to respond to cold exposure with increased thermogenesis, underscoring the importance of adiponectin for BAT function^23^. In summary, we found increased mRNA expression of three genes involved in nonshivering thermogenesis and whose function is regulated by the sympathetic nervous system (SNS). SNS dysfunction is a hallmark of obesity and insulin resistance states^24,25,26,27,28^. SNS dysfunction, as observed in obesity, is associated with low adiponectin levels^29^. SG and medical weight loss in humans have been shown to modulate SNS activity^30,31^. Although hyperleptinemia and hyperinsulinemia, in obesity, have been considered the drivers of SNS dysfunction, there is still considerable debate on this matter. Studies, so far, have found a weakly positive or no correlation between plasma leptin and plasma insulin and SNS activity in humans with obesity^32,33,34,35,36^. The finding of an equally reduced plasma insulin and leptin level in the SG group compared to SWM and SH groups, and SWM group, respectively, speaks against the reduction in those hormones as being associated with the changes in energy homeostasis observed in the SG group.

Glucose tolerance improved significantly in the SG group, to a similar magnitude as the SWM group. It is true, also, that glucose tolerance improved in all groups, relative to baseline. Since all the mice received enrofloxacin for 7 days after surgeries, it is intriguing whether microbiome changes could have contributed to the glucose tolerance improvement. Food intake cannot be ascribed as the main reason for improved glucose homeostasis, as the SG group consumed more calories than the SWM group but maintained a similar body weight. Considering the other findings in BAT, we wonder if increased metabolic activity in that fat depot was linked to the improvement in glucose homeostasis. It has been shown that BAT participates in glucose homeostasis, by promoting an increase in glucose uptake by metabolically active BAT and white adipose tissue, as observed in mice that received BAT transplants^37^. However, we do not know if our findings in BAT are connected to the SG improvement in blood glucose. The influence of sex hormones on metabolism is indisputable^38^. A decrease in estrogens during menopause is associated with increased fat mass accumulation and decreased lean mass^39,40^. We did not find evidence of ovarian failure in the mice in our study, as indicated by vaginal smears reflecting various phases of the estrus cycle.

Future study should focus on the many questions arising from this study’s findings, including the role of increased energy expenditure and increased fecal lipid excretion in the weight loss and blood glucose improvement observed in mature, obese mice that undergo SG.

## Conflict of interest statement

none

## Notes

### Competing Interest Statement

The authors have declared no competing interest.

## References

1. Caspard H, Jabbour S, Hammar N, Fenici P, Sheehan JJ, Kosiborod M. Recent trends in the prevalence of type 2 diabetes and the association with abdominal obesity lead to growing health disparities in the USA: An analysis of the NHANES surveys from 1999 to 2014. Diabetes Obes Metab. 2018 Mar;20(3):667–671. doi: 10.1111/dom.13143. Epub 2017 Dec 1. PMID: 29077244; PMCID: PMC5836923.

2. Arterburn D, Wellman R, Emiliano A, Smith SR, Odegaard AO, Murali S, Williams N, Coleman KJ, Courcoulas A, Coley RY, Anau J, Pardee R, Toh S, Janning C, Cook A, Sturtevant J, Horgan C, McTigue KM; PCORnet Bariatric Study Collaborative. Comparative Effectiveness and Safety of Bariatric Procedures for Weight Loss: A PCORnet Cohort Study. Ann Intern Med. 2018 Dec 4;169(11):741–750. doi: 10.7326/M17-2786. Epub 2018 Oct 30. PMID: 30383139; PMCID: PMC6652193.

3. Schauer PR, Kashyap SR, Wolski K, Brethauer SA, Kirwan JP, Pothier CE, Thomas S, Abood B, Nissen SE, Bhatt DL. Bariatric surgery versus intensive medical therapy in obese patients with diabetes. N Engl J Med. 2012 Apr 26;366(17):1567–76. doi: 10.1056/NEJMoa1200225. Epub 2012 Mar 26. PMID: 22449319; PMCID: PMC3372918.

4. Schauer PR, Bhatt DL, Kirwan JP, Wolski K, Brethauer SA, Navaneethan SD, Aminian A, Pothier CE, Kim ES, Nissen SE, Kashyap SR; STAMPEDE Investigators. Bariatric surgery versus intensive medical therapy for diabetes--3-year outcomes. N Engl J Med. 2014 May 22;370(21):2002–13. doi: 10.1056/NEJMoa1401329. Epub 2014 Mar 31. PMID: 24679060; PMCID: PMC5451259.

5. Schauer PR, Bhatt DL, Kirwan JP, Wolski K, Aminian A, Brethauer SA, Navaneethan SD, Singh RP, Pothier CE, Nissen SE, Kashyap SR; STAMPEDE Investigators. Bariatric Surgery versus Intensive Medical Therapy for Diabetes - 5-Year Outcomes. N Engl J Med. 2017 Feb 16;376(7):641–651. doi: 10.1056/NEJMoa1600869. PMID: 28199805; PMCID: PMC5451258.

6. English WJ, DeMaria EJ, Brethauer SA, Mattar SG, Rosenthal RJ, Morton JM. American Society for Metabolic and Bariatric Surgery estimation of metabolic and bariatric procedures performed in the United States in 2016. Surg Obes Relat Dis. 2018 Mar;14(3):259–263. doi: 10.1016/j.soard.2017.12.013. Epub 2017 Dec 16. PMID: 29370995.

7. Buchwald H, Avidor Y, Braunwald E, Jensen MD, Pories W, Fahrbach K, Schoelles K. Bariatric surgery: a systematic review and meta-analysis. JAMA. 2004 Oct 13;292(14):1724–37. doi: 10.1001/jama.292.14.1724. Erratum in: JAMA. 2005 Apr 13;293(14):1728. PMID: 15479938.

8. Courcoulas AP, King WC, Belle SH, Berk P, Flum DR, Garcia L, Gourash W, Horlick M, Mitchell JE, Pomp A, Pories WJ, Purnell JQ, Singh A, Spaniolas K, Thirlby R, Wolfe BM, Yanovski SZ. Seven-Year Weight Trajectories and Health Outcomes in the Longitudinal Assessment of Bariatric Surgery (LABS) Study. JAMA Surg. 2018 May 1;153(5):427–434. doi: 10.1001/jamasurg.2017.5025. PMID: 29214306; PMCID: PMC6584318.

9. Ryan KK, Tremaroli V, Clemmensen C, Kovatcheva-Datchary P, Myronovych A, Karns R, Wilson-Pérez HE, Sandoval DA, Kohli R, Bäckhed F, Seeley RJ. FXR is a molecular target for the effects of vertical sleeve gastrectomy. Nature. 2014 May 8;509(7499):183–8. doi: 10.1038/nature13135. Epub 2014 Mar 26. PMID: 24670636; PMCID: PMC4016120.

10. Saeidi N, Meoli L, Nestoridi E, Gupta NK, Kvas S, Kucharczyk J, Bonab AA, Fischman AJ, Yarmush ML, Stylopoulos N. Reprogramming of intestinal glucose metabolism and glycemic control in rats after gastric bypass. Science. 2013 Jul 26;341(6144):406–10. doi: 10.1126/science.1235103. PMID: 23888041; PMCID: PMC4068965.

11. McGavigan AK, Garibay D, Henseler ZM, Chen J, Bettaieb A, Haj FG, Ley RE, Chouinard ML, Cummings BP. TGR5 contributes to glucoregulatory improvements after vertical sleeve gastrectomy in mice. Gut. 2017 Feb;66(2):226–234. doi: 10.1136/gutjnl-2015-309871. Epub 2015 Oct 28. PMID: 26511794; PMCID: PMC5512436.

12. Griffin C, Hutch CR, Abrishami S, Stelmak D, Eter L, Li Z, Chang E, Agarwal D, Zamarron B, Varghese M, Subbaiah P, MacDougald OA, Sandoval DA, Singer K. Inflammatory responses to dietary and surgical weight loss in male and female mice. Biol Sex Differ. 2019 Apr 3;10(1):16. doi: 10.1186/s13293-019-0229-7. PMID: 30944030; PMCID: PMC6446331.

13. Frohman HA, Rychahou PG, Li J, Gan T, Evers BM. Development of murine bariatric surgery models: lessons learned. J Surg Res. 2018 Sep;229:302–310. doi: 10.1016/j.jss.2018.04.022. Epub 2018 May 10. PMID: 29937006; PMCID: PMC6298430.

14. Champy MF, Selloum M, Zeitler V, Caradec C, Jung B, Rousseau S, Pouilly L, Sorg T, Auwerx J. Genetic background determines metabolic phenotypes in the mouse. Mamm Genome. 2008 May;19(5):318–31. doi: 10.1007/s00335-008-9107-z. Epub 2008 Apr 5. PMID: 18392653.

15. Garibay D, Cummings BP. A Murine Model of Vertical Sleeve Gastrectomy. J Vis Exp. 2017 Dec 18;(130):56534. doi: 10.3791/56534. PMID: 29286478; PMCID: PMC5755614.

16. Kiess W, Galler A, Reich A, Müller G, Kapellen T, Deutscher J, Raile K, Kratzsch J. Clinical aspects of obesity in childhood and adolescence. Obes Rev. 2001 Feb;2(1):29–36. doi: 10.1046/j.1467-789x.2001.00017.x. PMID: 12119634.

17. Lupoli R, Lembo E, Saldalamacchia G, Avola CK, Angrisani L, Capaldo B. Bariatric surgery and long-term nutritional issues. World J Diabetes. 2017 Nov 15;8(11):464–474. doi: 10.4239/wjd.v8.i11.464. PMID: 29204255; PMCID: PMC5700383.

18. Bäckhed F, Ding H, Wang T, Hooper LV, Koh GY, Nagy A, Semenkovich CF, Gordon JI. The gut microbiota as an environmental factor that regulates fat storage. Proc Natl Acad Sci U S A. 2004 Nov 2;101(44):15718–23. doi: 10.1073/pnas.0407076101. Epub 2004 Oct 25. PMID: 15505215; PMCID: PMC524219.

19. Crichton PG, Lee Y, Kunji ER. The molecular features of uncoupling protein 1 support a conventional mitochondrial carrier-like mechanism. Biochimie. 2017 Mar;134:35–50. doi: 10.1016/j.biochi.2016.12.016. Epub 2017 Jan 3. PMID: 28057583; PMCID: PMC5395090.

20. Feldmann HM, Golozoubova V, Cannon B, Nedergaard J. UCP1 ablation induces obesity and abolishes diet-induced thermogenesis in mice exempt from thermal stress by living at thermoneutrality. Cell Metab. 2009 Feb;9(2):203–9. doi: 10.1016/j.cmet.2008.12.014. PMID: 19187776.

21. Sell H, Berger JP, Samson P, Castriota G, Lalonde J, Deshaies Y, Richard D. Peroxisome proliferator-activated receptor gamma agonism increases the capacity for sympathetically mediated thermogenesis in lean and ob/ob mice. Endocrinology. 2004 Aug;145(8):3925–34. doi: 10.1210/en.2004-0321. Epub 2004 May 6. PMID: 15131020.

22. Puigserver P, Spiegelman BM. Peroxisome proliferator-activated receptor-gamma coactivator 1 alpha (PGC-1 alpha): transcriptional coactivator and metabolic regulator. Endocr Rev. 2003 Feb;24(1):78–90. doi: 10.1210/er.2002-0012. PMID: 12588810.

23. Wei Q, Lee JH, Wang H, Bongmba OYN, Wu CS, Pradhan G, Sun Z, Chew L, Bajaj M, Chan L, Chapkin RS, Chen MH, Sun Y. Adiponectin is required for maintaining normal body temperature in a cold environment. BMC Physiol. 2017 Oct 23;17(1):8. doi: 10.1186/s12899-017-0034-7. PMID: 29058611; PMCID: PMC5651620.

24. Grassi G, Seravalle G, Dell’Oro R, Turri C, Bolla GB, Mancia G. Adrenergic and reflex abnormalities in obesity-related hypertension. Hypertension. 2000 Oct;36(4):538–42. doi: 10.1161/01.hyp.36.4.538. PMID: 11040232.

25. Alvarez GE, Beske SD, Ballard TP, Davy KP. Sympathetic neural activation in visceral obesity. Circulation. 2002 Nov 12;106(20):2533–6. doi: 10.1161/01.cir.0000041244.79165.25. PMID: 12427647.

26. Rumantir MS, Vaz M, Jennings GL, Collier G, Kaye DM, Seals DR, Wiesner GH, Brunner-La Rocca HP, Esler MD. Neural mechanisms in human obesity-related hypertension. J Hypertens. 1999 Aug;17(8):1125–33. doi: 10.1097/00004872-199917080-00012. PMID: 10466468.

27. Lee ZS, Critchley JA, Tomlinson B, Young RP, Thomas GN, Cockram CS, Chan TY, Chan JC. Urinary epinephrine and norepinephrine interrelations with obesity, insulin, and the metabolic syndrome in Hong Kong Chinese. Metabolism. 2001 Feb;50(2):135–43. doi: 10.1053/meta.2001.19502. PMID: 11229419.

28. Huggett RJ, Hogarth AJ, Mackintosh AF, Mary DA. Sympathetic nerve hyperactivity in non-diabetic offspring of patients with type 2 diabetes mellitus. Diabetologia. 2006 Nov;49(11):2741–4. doi: 10.1007/s00125-006-0399-9. Epub 2006 Sep 13. PMID: 16969648.

29. Nowak Ł, Adamczak M, Wiecek A. Blockade of sympathetic nervous system activity by rilmenidine increases plasma adiponectin concentration in patients with essential hypertension. Am J Hypertens. 2005 Nov;18(11):1470–5. doi: 10.1016/j.amjhyper.2005.05.026. PMID: 16280284.

30. Mark AL, Norris AW, Rahmouni K. Sympathetic inhibition after bariatric surgery. Hypertension. 2014 Aug;64(2):235–6. doi: 10.1161/HYPERTENSIONAHA.114.03183. PMID: 24866141; PMCID: PMC4184982.

31. Seravalle G, Colombo M, Perego P, Giardini V, Volpe M, Dell’Oro R, Mancia G, Grassi G. Long-term sympathoinhibitory effects of surgically induced weight loss in severe obese patients. Hypertension. 2014 Aug;64(2):431–7. doi: 10.1161/HYPERTENSIONAHA.113.02988. Epub 2014 May 27. PMID: 24866140. Stanford KI, Middelbeek RJ, Townsend KL, An D, Nygaard EB, Hitchcox KM, Markan KR, Nakano K, Hirshman MF, Tseng YH, Goodyear LJ. Brown adipose tissue regulates glucose homeostasis and insulin sensitivity. J Clin Invest. 2013 Jan;123(1):215–23. doi: 10.1172/JCI62308. Epub 2012 Dec 10. PMID: 23221344; PMCID: PMC3533266.

32. Vollenweider P, Randin D, Tappy L, Jéquier E, Nicod P, Scherrer U. Impaired insulin-induced sympathetic neural activation and vasodilation in skeletal muscle in obese humans. J Clin Invest. 1994 Jun;93(6):2365–71. doi: 10.1172/JCI117242. PMID: 8200969; PMCID: PMC294442.

33. Straznicky NE, Lambert GW, Masuo K, Dawood T, Eikelis N, Nestel PJ, McGrane MT, Mariani JA, Socratous F, Chopra R, Esler MD, Schlaich MP, Lambert EA. Blunted sympathetic neural response to oral glucose in obese subjects with the insulin-resistant metabolic syndrome. Am J Clin Nutr. 2009 Jan;89(1):27–36. doi: 10.3945/ajcn.2008.26299. Epub 2008 Dec 3. PMID: 19056585.

34. Snitker S, Pratley RE, Nicolson M, Tataranni PA, Ravussin E. Relationship between muscle sympathetic nerve activity and plasma leptin concentration. Obes Res. 1997 Jul;5(4):338–40. doi: 10.1002/j.1550-8528.1997.tb00561.x. PMID: 9285841.

35. Monroe MB, Van Pelt RE, Schiller BC, Seals DR, Jones PP. Relation of leptin and insulin to adiposity-associated elevations in sympathetic activity with age in humans. Int J Obes Relat Metab Disord. 2000 Sep;24(9):1183–7. doi: 10.1038/sj.ijo.0801364. PMID: 11033988.

36. Narkiewicz K, Kato M, Phillips BG, Pesek CA, Choe I, Winnicki M, Palatini P, Sivitz WI, Somers VK. Leptin interacts with heart rate but not sympathetic nerve traffic in healthy male subjects. J Hypertens. 2001 Jun;19(6):1089–94. doi: 10.1097/00004872-200106000-00014. PMID: 11403358.

37. Stanford KI, Middelbeek RJ, Townsend KL, An D, Nygaard EB, Hitchcox KM, Markan KR, Nakano K, Hirshman MF, Tseng YH, Goodyear LJ. Brown adipose tissue regulates glucose homeostasis and insulin sensitivity. J Clin Invest. 2013 Jan;123(1):215–23. doi: 10.1172/JCI62308. Epub 2012 Dec 10. PMID: 23221344; PMCID: 33266.

38. Leeners B, Geary N, Tobler PN, Asarian L. Ovarian hormones and obesity. Hum Reprod Update. 2017 May 1;23(3):300–321. doi: 10.1093/humupd/dmw045. PMID: 28333235; PMCID: PMC5850121.

39. Kanaley JA, Sames C, Swisher L, Swick AG, Ploutz-Snyder LL, Steppan CM, Sagendorf KS, Feiglin D, Jaynes EB, Meyer RA, Weinstock RS. Abdominal fat distribution in pre- and postmenopausal women: The impact of physical activity, age, and menopausal status. Metabolism. 2001 Aug;50(8):976–82. doi: 10.1053/meta.2001.24931. PMID: 11474488.

40. Franklin RM, Ploutz-Snyder L, Kanaley JA. Longitudinal changes in abdominal fat distribution with menopause. Metabolism. 2009 Mar;58(3):311–5. doi: 10.1016/j.metabol.2008.09.030. PMID: 19217444.

